# Regenotyping structural variants through an accurate force-calling method

**DOI:** 10.1101/2022.08.29.505534

**Authors:** Tao Jiang, Shuqi Cao, Yadong Liu, Shiqi Liu, Bo Liu, GuoHua Wang, Yadong Wang

## Abstract

Long-read sequencing technologies have great potential for the comprehensive discovery of structural variation (SV). However, accurate genotype assignment for SV is still a challenge due to unavoidable factors, such as specific sequencing errors or limited coverage. Herein, we propose cuteSV2, a fast and accurate long-read-based regenotyping approach that is used to force calling genotypes for given records. In cuteSV2, which is an upgraded version of cuteSV, an improved refinement strategy is applied on the signatures, and the heuristic extracted signatures are purified through spatial and allele similarity estimation. The benchmarking results on several baseline evaluations demonstrate that cuteSV2 outperforms the state-of-the-art methods and is a scalable and robust approach for population studies and clinical practice. cuteSV2 is available at https://github.com/tjiangHIT/cuteSV.

## Introduction

The structural variant (SV) is a fundamental type of genomic mutation that is strongly associated with evolution, population structure, and clinical diseases^[1-3]^. Typically, an SV is larger than 50 bp in size and mainly contains insertions, deletions, inversions, duplications, translocations, and more complex variants^[4]^. Previous population studies, such as genome-wide association studies (GWAS), focused on the exhaustive characterization of single-nucleotide variants (SNVs) underlying human traits and diseases and ignored the important heritability of SVs^[5, 6]^. However, because of the variety and wide-range alterations of SVs, a large number of nucleotides in the genome are usually affected, which is of significant value in genome research. Recently, long-read sequencing technologies have enabled the discovery of full-spectrum SVs on a single individual^[7-10]^, which provides an opportunity to reveal population-based SVs through large-scale sequencing^[11-14]^.

Currently, the joint variant calling strategy is often applied for the construction of population-based SV genetic maps. All samples can be considered simultaneously, and high-quality population-scale variant callsets, which can provide the overall variant distribution of the population, can be generated. To achieve better joint calling performance, the critical step is to perform regenotyping for each sample at all SV positions to obtain an accurate allele frequency distribution in the population. In the regenotyping of variations, the original SV genotypes are refined or filled due to insufficient coverage^[15, 16]^. Compared with short-read sequencing, in long-read sequencing, such as Pacific Bioscience (PacBio) and Oxford Nanopore Technology (ONT), longer reads that can cross whole variation regions can be generated. However, unignorable sequencing errors and high expenses restrict its wide application in SV genetic studies and make it difficult to obtain reliable genotypes for a given SV across a population group.

Many force calling methods have been developed to achieve accurate SV regenotyping with long-read sequencing technologies. SVJedi^[17]^ is used to filter out the informative alignments to quantify the presence of SV alleles. However, it still lacks high accuracy and only accepts the original reads as input, which requires a large amount of time to accomplish read alignment. In another widely used method, Sniffles1^[18]^, a self-balancing tree is constructed to compute the fraction of reads, and the genotype is considered based on the allele frequency. However, it focuses on SV detection and always makes mistakes in determining genotypes for SVs. In Sniffles2^[19]^, which is a redesigned version of Sniffles1, a three-phase clustering process is employed to achieve better performance in genotyping. In general, such force calling methods have drawbacks in determining the real correct genotype for SVs. The inaccuracy and inconvenience of regenotyping are still bottlenecks in the construction of highly resolved population-based SV genetics maps.

Here, we introduce an accurate and fast regenotyping method that uses force calling, named cuteSV2, to determine the zygosities of all SVs on each individual at the population scale. Using a bisection search and quasi-Markov model-based clustering strategy on the diverse alignment signatures, the zygosities are obtained by calculating the read distribution likelihood of each candidate circumstance. Additionally, a scanning line is designed in cuteSV2 to decrease the elapsed time of read hunting and increase whole force calling. The experiments are implemented on an Ashkenazim trio-family and a population group with 100 Chinese individuals. The results indicate that our regenotyping method cuteSV2 has an outstanding and a reliable performance compared to homogeneous state-of-the-art methods, which indicates that it will assist in obtaining accurate allele frequencies of homology SVs for further population genetic measurement and estimation.

## Results

### 1. Overview of cuteSV2

In cuteSV2, the individual sorted BAM files and reference genome and regenotyped VCF files of target SVs are taken as input to achieve the genotype force calling of all regenotyped SVs on the individual. The four major steps developed in cuteSV2 for accurate and fast force calling are given as follows:

#### Step 1

Apply the multiple signature extraction module in cuteSV to comprehensively collect the signatures of various types of SVs.

#### Step 2

Mark all candidate signatures for each given SV through a specifically designed spatial similarity measurement and allele similarity estimation.

#### Step 3

Overlap alignment reads with given SVs via a scanning line on the chromosome to obtain the read distribution around the SVs.

#### Step 4

Apply the likelihood estimation module in cuteSV to reassign the genotype for each given SV on the corresponding sample.

A schematic illustration is shown in Fig. 1, and more details are provided in the “Methods” section.

**Fig. 1.**
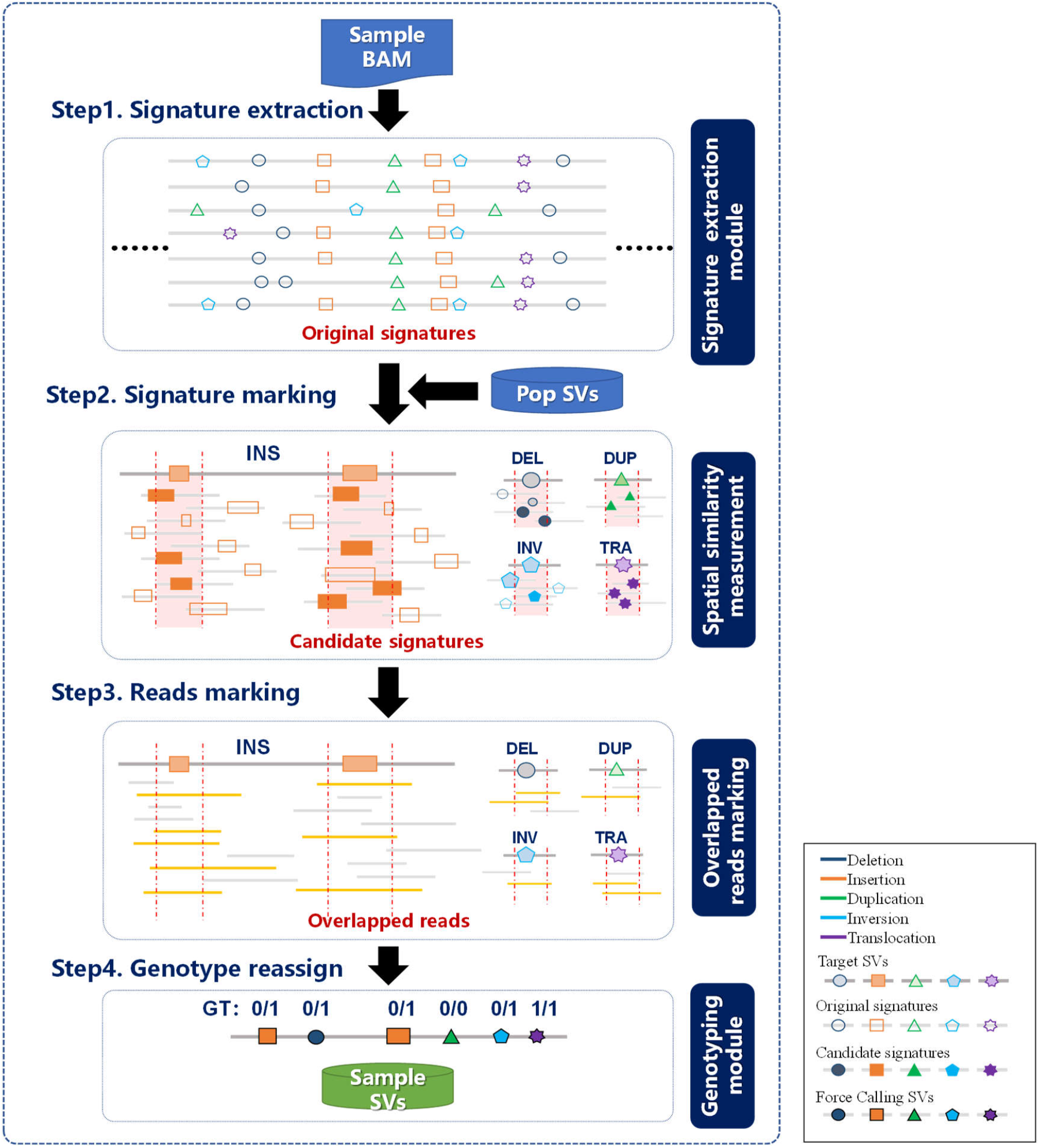
Schematic illustration of the cuteSV2 approach. **Step 1**. The signature extraction module is applied to comprehensively collect the signatures of various types of SVs. **Step 2**. All candidate signatures for each given SV are marked through specifically designed spatial similarity measurements and allele similarity estimation. **Step 3**. The overlapping alignment reads with given SVs are marked using the *scanning line* algorithm to acquire the detailed distribution of reads around SVs. **Step 4**. The genotype for each given SV on the corresponding sample is reassigned through maximum likelihood estimation.

The key points of cuteSV2 are the heuristic statistics of the alignment reads and variation signatures, which enable the accurate genotype assignment. In step 1 of cuteSV2, the signature extraction module of cuteSV is inherited with several further prunings to collect comprehensive variation signatures and remove the noise from the sequencing and alignment procedure, thus providing a solid foundation for subsequent processing. Then, in step 2 and step 3 of cuteSV2, the specially designed algorithms measure the spatial similarity and allele similarity between signatures and target SVs and collect the candidate signatures heuristically, yielding precise read distribution statistics and the correct likelihood estimation of zygosities.

In addition, the large-scale population group has massive data for processing and analyzing, placing extreme pressure on computational and storage resources. In step 1 of cuteSV2, the external disk is used for recording the temporary files so that the memory footprint of the regenotyping method is greatly reduced. Furthermore, a block divide-and-conquer approach is applied to process force calling in parallel with multiple CPU cores. The effective implementation greatly speeds up cuteSV2 under various conditions in practice. More details are shown below.

### 2. Benchmarks of regenotyping performance on HG002 datasets

#### 2.1 Evaluations on different sequencing platforms

We first evaluated the regenotyping performance of cuteSV2 and other state-of-the-art methods (i.e., Sniffles1^[18]^, Sniffles2^[19]^, and SVJedi^[17]^) on a well-known human sample HG002 from the Ashkenazim trio-family group based on the Genome in a Bottle (GiaB) ground truth set (SV v0.6) from the National Institute of Standards and Technology (NIST). The target SV candidates were integrated from the Ashkenazim trio-family group (i.e., HG002, HG003, and HG004), as detected by PBSV (https://github.com/PacificBiosciences/pbsv). The benchmarking results (Fig. 2A and Supplementary Tables S1∼3) indicate that the highest precision (96.80% for HiFi, 96.50% for CLR, 96.99% for ONT) and the fewest false-positive SVs among the four methods were achieved using cuteSV2, while approximately twice as many false-positive SVs and lower precision were observed for the other three methods reported. In regard to the recall rate, cuteSV2, Sniffles1, and Sniffles2 had competitive performance against the other methods, in addition to SVJedi. More specifically, the highest recall on HiFi (92.71%), the second highest on CLR (94.34% compared with 94.63% of Sniffles1), and the second highest on ONT (94.03% compared with 94.26% of Sniffles2), were achieved using cuteSV2, which proves its high and stable recall rate. Overall, the best F1 scores under the diverse sequencing technologies (94.71% for HiFi, 95.41% for CLR, 95.49% for ONT), were obtained using cuteSV2, proving its high precision and stable recall rate. Furthermore, we assessed the genotype concordance between the regenotyped results and the ground truth. It is obvious that the best concordance (98.21% for HiFi, 97.8% for CLR, 98.2% for ONT) was obtained by cuteSV2, followed by Sniffles2 (97.12% for HiFi, 97.74% for CLR, 96.53% for ONT), while the other two methods performed relatively worse (Sniffles1: 46.20% for HiFi, 46.13% for CLR, 94.49% for ONT, SVJedi: 71.58% for HiFi, 91.36% for CLR, and 87.13% for ONT).

**Fig. 2.**
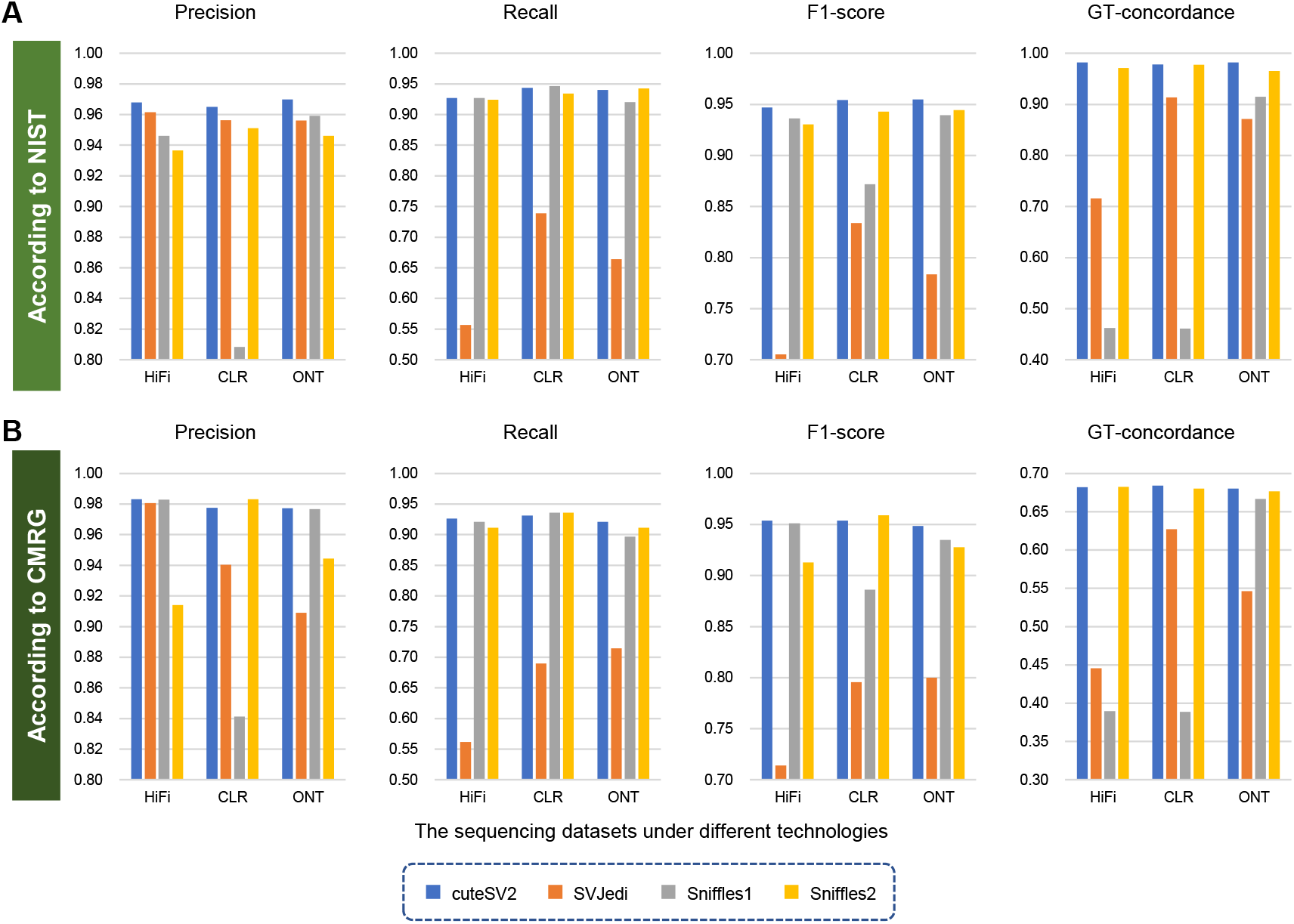
The benchmark results under various sequencing datasets on the HG002 sample. **A**. The precision, recall, F1 score, and genotype concordance under different sequencing technologies, including HiFi (PacBio HiFi sequencing), CLR (PacBio CLR sequencing), and ONT (Oxford Nanopore Technologies sequencing). The ground truth comes from the Genome in a Bottle (GiaB) ground truth set (SV v0.6) from the National Institute of Standards and Technology (NIST). **B**. The precision, recall, F1 score, and genotype concordance under different sequencing technologies, including HiFi, CLR, and ONT. The ground truth comes from the GIAB Challenging Medically Relevant Gene Benchmark (CMRG) v1.00.

We then evaluated the performance of regenotyping on the GIAB Challenging Medically Relevant Gene Benchmark (CMRG) v1.00 (Fig. 2B and Supplementary Tables S4∼6). For precision and recall, cuteSV2 was the best among all methods regardless of the sequencing technologies, and the highest precision and recall on the HiFi and ONT datasets and the second highest on the CLR datasets were obtained using cuteSV2, whose performance was slightly behind Sniffles2. Considering the sensitivity and accuracy at the same time, the highest F1 scores on HiFi and ONT data and the second best on CLR data (0.53% lower than that of Sniffles2) were obtained using cuteSV2. In regard to the genotype concordance, cuteSV2 and Sniffles2 performed much better than the other two methods, while there was no more than a 1% gap between these two methods.

#### 2.2 Evaluations of different sequencing depths

Furthermore, we randomly downsampled the origin HG002 datasets to 30×, 20×, 10× and 5× and regenotyped HG002 on them (Supplementary Tables S1∼6). When the sequencing coverage decreased, the performance of all the methods decreased to some extent (Fig. 3A); however, the best performance in most conditions was achieved using cuteSV2. Especially focusing on precision, the performance of cuteSV2 only decreased by less than 1% and even slightly increased when the coverage varied from the origin coverage to 5× (from 96.8% to 97.05% for HiFi, from 96.5% to 95.35% for CLR, and from 96.99% to 96.97% for ONT) and was higher than that of the other methods. It is worth mentioning that SVJedi showed an extremely high precision under low coverage conditions. However, its high precision was due to the few false-positives with few true positives, and less than one-fifth true positive SVs were obtained using SVJedi, demonstrating its extremely insufficient performance. The recall rate of each method decreased more because of the absence of pivotal SV signatures in low-coverage data. The performance of cuteSV2 and Sniffles2 dropped by approximately 6% when the coverage decreased to 10×, Sniffles1 dropped by approximately 9% and SVJedi dropped much more at over 20%. When the coverage was decreased to 5×, the recall rate had a large gap: cuteSV2 and Sniffles2 fell over 10%, and Sniffles1 and SVJedi fell more, especially on the CLR dataset. Therefore, it is reasonable to draw the conclusion that sequencing with at least 10× is needed for accurate regenotyping, and lower coverage may lead to insufficient genotyping. Comprehensively considering precision and recall, the best F1 scores on various coverage data under all sequencing technologies were still achieved using cuteSV2. In regard to genotype concordance, the best performance on low coverage datasets (dropped by 8.24% for HiFi, 15.48% for CLR, 12.98% for ONT) was obtained using Sniffles2, with cuteSV2 following close (dropped by 12.37% for HiFi, 18.57% for CLR, 14.13% for ONT).

**Fig. 3.**
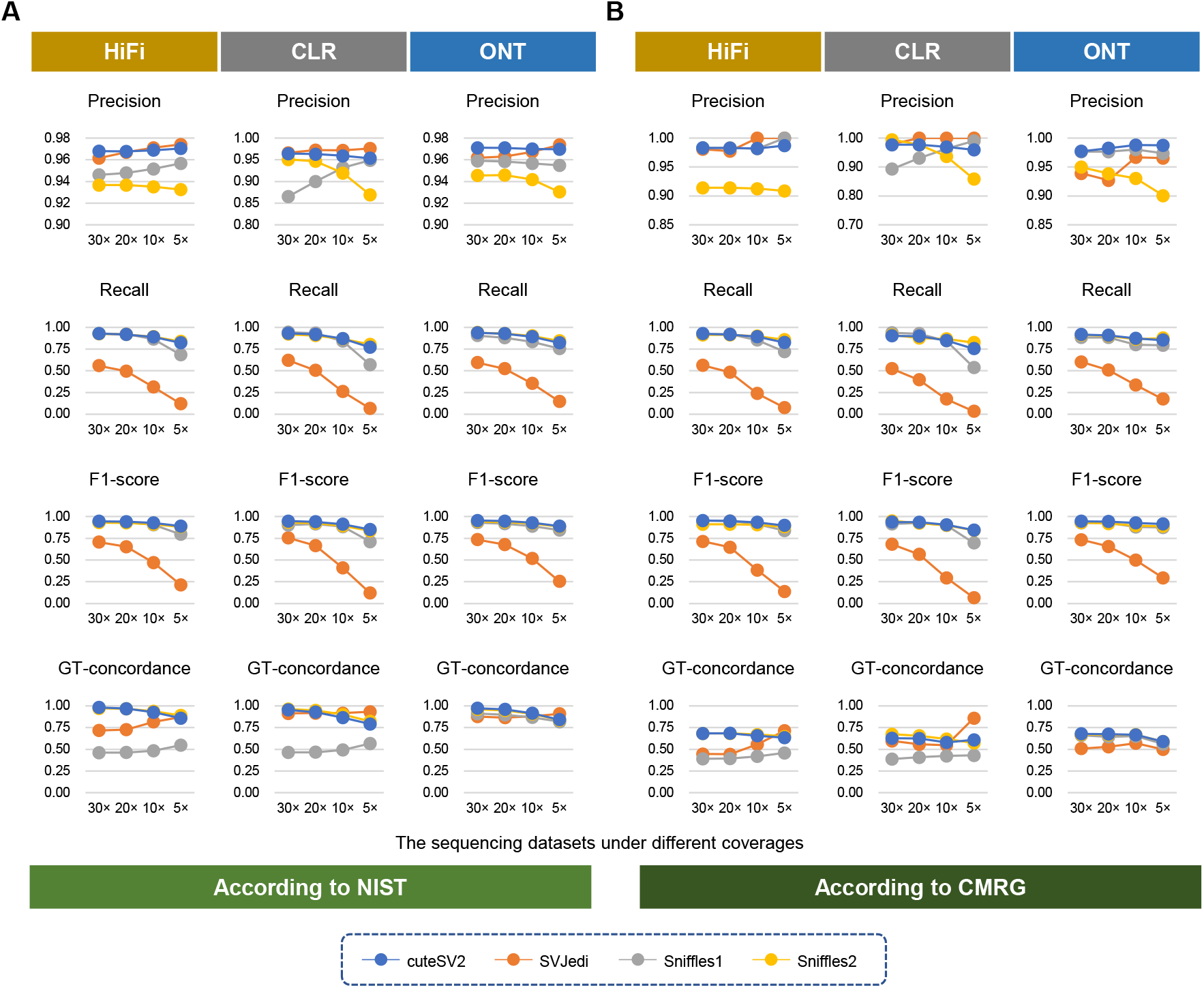
The benchmark results under various sequencing depths on the HG002 sample. **A**. The precision, recall, F1 score, and genotype concordance under different sequencing coverages (i.e., 30×, 20×, 10×, and 5×) using different sequencing technologies (i.e., HiFi, CLR, and ONT) based on the NIST ground truth. **B**. The precision, recall, F1 score, and genotype concordance under different sequencing coverages (i.e., 30×, 20×, 10×, and 5×) on different sequencing technologies (i.e., HiFi, CLR, and ONT) based on the CMRG ground truth.

We also evaluated the performance on low-coverage datasets of the CMRG ground truth, and the benchmark results showed a similar tendency as above (Fig. 3B). A high and stable precision was observed when using cuteSV2, Sniffles1 and SVJedi; however, an apparent decrease in precision was observed for Sniffles2. In terms of the recall rate, a similar decrease was observed for cuteSV2, Sniffles1, and Sniffles2. Moreover, the performance of SVJedi was always much worse, especially in low coverage. Therefore, the highest F1 score under various sequencing coverages was still obtained using cuteSV2. Meanwhile, better and closer genotype concordance was obtained using cuteSV2 and Sniffles2 compared with the other two methods, and compared with each other, cuteSV2 performed better on the PacBio HiFi and ONT sequencing datasets, while Sniffles2 performed better on PacBio CLR sequencing.

### 3. Benchmarks of regenotyping performance on a large-scale Chinese cohort

#### 3.1 Genetic testing of genotyping results

To evaluate the regenotyping performance on a large-scale cohort, we applied cuteSV2, Sniffles1, and Sniffles2 (SVJedi was discarded due to its relatively low performance) on a real population group consisting of 100 Chinese individuals. Data from these individuals were sequenced using ONT technology with approximately 10∼15 ×, and the target SVs were achieved using a population SV calling pipeline including SV discovery and integration (see *Methods* for more details). After regenotyping, Bcftools^[20]^ was adopted to calculate the allele frequency (AF), the test score of the Hardy–Weinberg equilibrium (HWE), and the test score of excess heterozygosity (ExcHet) for each variant. After removing the nonexistent SVs, whose AF was equal to 0, the largest number of SVs were retained using cuteSV2 (96,431 SVs for cuteSV2, 95,184 SVs for Sniffles2, and 73,918 for Sniffles1), which indicates the ability of cuteSV2 to detect abundant SVs in the population (Supplementary Table S7). Meanwhile, we grouped the SVs by the allele frequency, and the distribution of SVs under different allele frequency intervals is shown in Fig. 4A. The distributions of cuteSV2 and Sniffles2 showed a similar tendency, while the distribution of Sniffles1 seemed imbalanced and had only a few SVs with allele frequencies larger than 0.5. In the intervals of low allele frequency, more SVs were obtained using cuteSV2, and in contrast, more SVs with high allele frequency were obtained using Sniffles2. It is worth mentioning that the largest number of singletons and doubletons (22,042 and 7,561, respectively) were recognized using cuteSV2, which is vital for the analysis of rare variants.

**Fig. 4.**
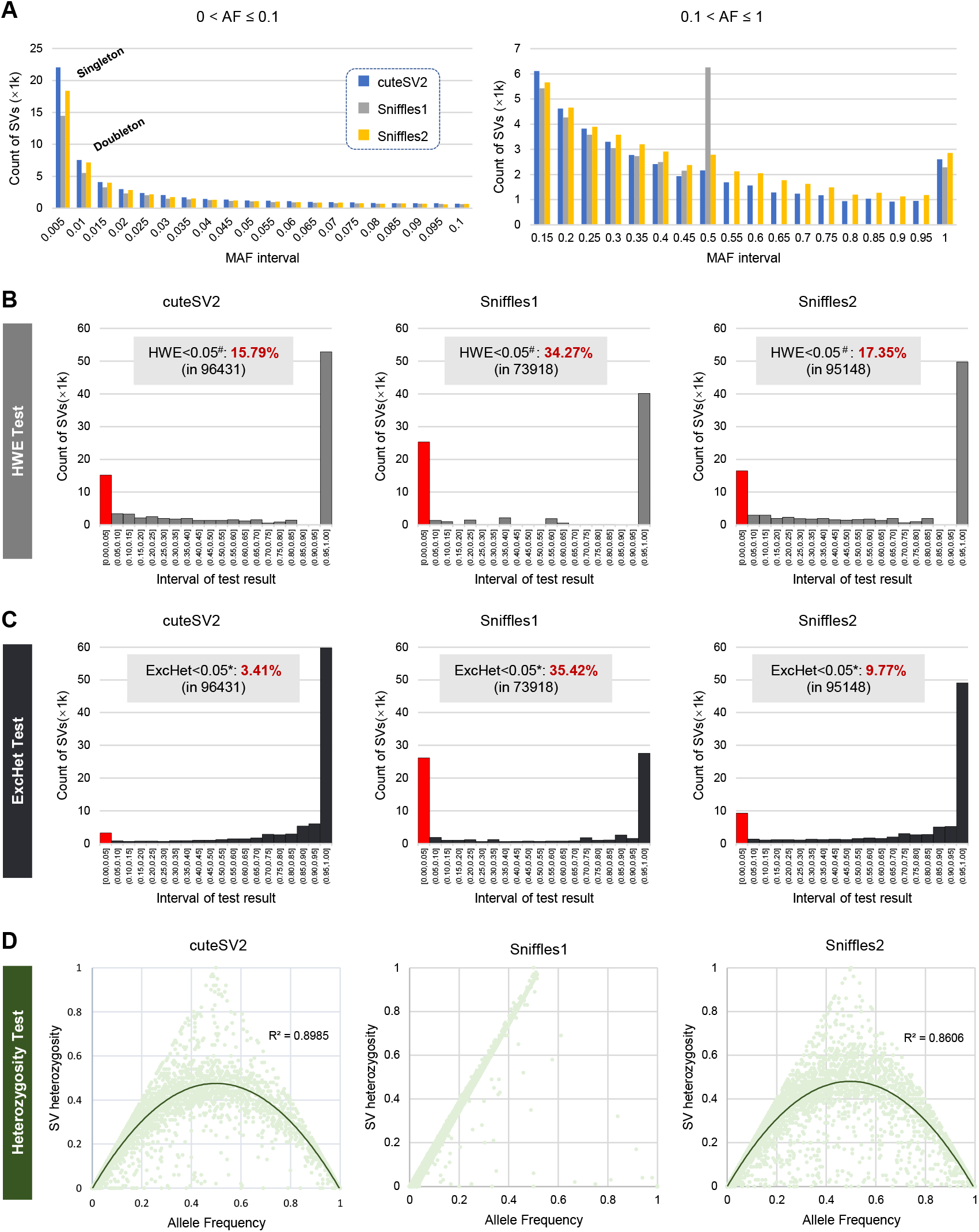
The benchmark genotyping results on the Chinese population group. **A**. The distribution of SV amounts under different minor allele frequency (MAF) intervals. The left panel shows the distribution of SVs whose MAF intervals are smaller than 0.1, and the right panel shows more common SVs, with MAF intervals larger than 0.1. It is also worth mentioning that the SVs in the regions of MAF=0.005 and MAF=0.01 indicate singletons and doubletons, respectively. **B**. The distribution of the Hardy–Weinberg equilibrium (HWE) test scores for these three methods in the population group. The red column and the label “HWE<0.05^#^” indicate low-quality SVs with missing rates higher than 5% or test scores lower than 0.05 and demonstrate HWE test failures. **C**. The distribution of test scores of excess heterozygosity for these three methods in the population group. The red column and the label “ExcHet<0.05*” indicate low-quality SVs with missing rates higher than 5% or test scores lower than 0.05 and demonstrate ExcHet test failures. **D**. The distribution of heterozygosity and allele frequency among 100 samples on chromosome 1. The line chart indicates the fitting curve of the distribution.

Then, we drew the distribution of the HWE test scores and ExcHet test scores of these three methods (Fig. 4B, 4C and Supplementary Tables S8-10) to evaluate the quality of population SV callsets from various methods. We first marked the SVs of low quality in the generated population SV callsets. The low-quality SVs consisted of two main parts. One part contained SVs with a missing rate larger than 5%, which indicated that over 5% of the SV alleles were missing, leading to uncertain SVs. The other part contained the SVs whose test scores were lower than 0.05, which indicated that these SVs had little agreement with population genetic regulation. As illustrated in Fig. 4B and 4C, in both tests, the lowest number of low-quality SVs together with the largest number of SVs with high scores (>0.95) were obtained using cuteSV2. In the HWE test, the lowest proportion of low-quality SVs (15.79%) was obtained using cuteSV2, followed by Sniffles2 (17.35%) and Sniffles1 (34.27%). In the ExcHet test, only 3.41% of SVs of cuteSV2 were recorded as low-quality SVs, indicating its outstanding genotype assignments for heterozygosity, while more bad cases were observed using Sniffles2 and Sniffles1 (9.77% and 35.42%, respectively).

We also analyzed the heterozygosity distribution together with the allele frequency (Fig. 4D). The distributions obtained using cuteSV2 and Sniffles2 fit the curve of HWE regulation well. However, an unreasonable fit was observed for the distribution obtained using Sniffles1 because most SVs located in the left part had a low allele frequency, and the straight fitting line was incongruent with the HWE regulation. We further measured the R^2^ value of the fitting curve to better assess the fitting performance. The R^2^ value obtained using cuteSV2 (0.8985) was larger than that using Sniffles2 (0.8606), indicating the higher relation between the heterozygosity distribution of cuteSV2 and the HWE theoretical distribution.

#### 3.2 Allele frequency concordance analysis according to the worldwide cohort

Furthermore, we evaluated the above Chinese population SV callsets on a worldwide well-studied population group consisting of 64 haplotypes^[14]^ because these haplotypes were generated from genome assembly and were of rather high quality and reliability. We selected the SVs that were shared in the 64 haplotypes and the data from the 100 Chinese individuals and calculated the allele frequency discordance of the shared SVs (Supplementary Table S11). From Fig. 5A, most shared SVs had adjacent and consistent allele frequencies, and few SVs had a large gap in allele frequency, indicating the wide sharing of mutual SVs in the human population. Among the three methods, the most consistent SVs, whose allele frequency discordance was smaller than 0.05, were obtained using cuteSV2 (14,149 for cuteSV2, 10,964 for Sniffles1, and 12,849 for Sniffles2). We also measured the proportion of insertions and deletions in the shared SVs. The results shown in the line chart of Fig. 5A demonstrate that there was a greater occurrence of insertions than deletions in each method, and the line fluctuation of cuteSV2 was most stable among intervals, indicating a better distribution of SV callsets from cuteSV2.

**Fig. 5.**
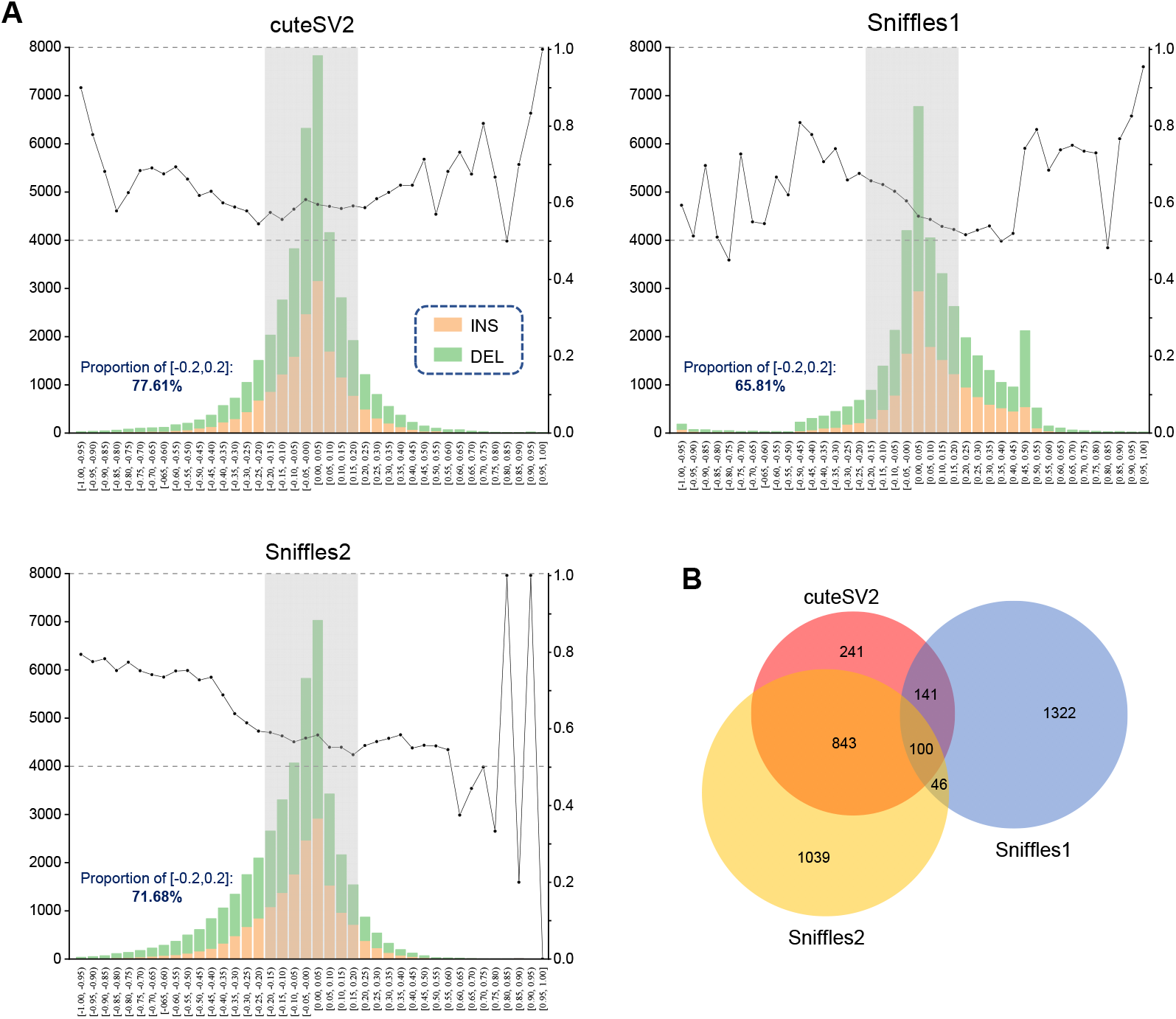
The benchmark results of the allele frequency concordance in the Chinese population group. **A**. The histogram represents the distribution of the allele frequency discordance between 64 well-studied haplotypes and our Chinese population SV callsets (that is, the difference in the allele frequency of the SVs that are shared in the worldwide cohort and our Chinese callsets). The line represents the amount distribution of different SV types, which is measured as the proportion of insertions divided by the amount of insertions (orange) and deletions (green). **B**. The Venn plot of the Chinese population SVs whose allele frequency discordance is larger than 0.5 for each SV caller.

Moreover, we analyzed the SVs that had a large allele frequency discordance (>0.5) with assembly haplotypes because these SVs may come from the erroneous recognition of methods, but they were also probably caused by genetic divergence in various population groups. The Venn plot in Fig. 5B indicates the sharing of these AF-discordant SVs among the three methods. The overlapping parts represent the discordant SVs reported by multiple methods, and the remaining parts indicate the unique discordant SVs of a single method. The smallest number of total discordant SVs were observed using cuteSV2 (1,325 for cuteSV2, 1,609 for Sniffles1, and 2,028 for Sniffles2), and most discordant SVs in cuteSV2 also appeared in Sniffles1 and Sniffles2. In other words, the fewest unique discordant SVs were reported using cuteSV2 (241 for cuteSV2, 1,322 for Sniffles1, and 1,039 for Sniffles2), which had the best concordance with the worldwide well-studied population group.

#### 3.3 Orthogonal validation of PacBio HiFi sequencing

We randomly selected three samples (Samples #1, #2, and #3) from this large-scale group to further assess the consistency of SV genotyping through validation using orthogonal PacBio HiFi sequencing. The SVs of low quality (defined above) were filtered, and only the remaining SVs were used for validation. From Fig. 6A, it is obvious that cuteSV2 and Sniffles2 showed an outstanding ability to genotype SVs, as over 14,000 remaining SVs for each sample were reported (14,387 for cuteSV2 and 16,352 for Sniffles2, on average), while approximately 6,686 SVs were obtained using Sniffles1 (Fig. 6A and Supplementary Tables S12∼14). In regard to genotype concordance, cuteSV2 and Sniffles2 were apparently better than Sniffles1, and the concordance of all AF intervals was higher than 90%. In terms of mass, the performances of cuteSV2 and Sniffles2 were approximate, as cuteSV2 had better concordance in higher AF intervals, and by contrast, Sniffles2 had better concordance in lower AF intervals. Taking the whole AF intervals into account, the highest genotype concordance was obtained using cuteSV2 (94.62% for cuteSV2, 21.90% for Sniffles1, and 94.11% for Sniffles2).

**Fig. 6.**
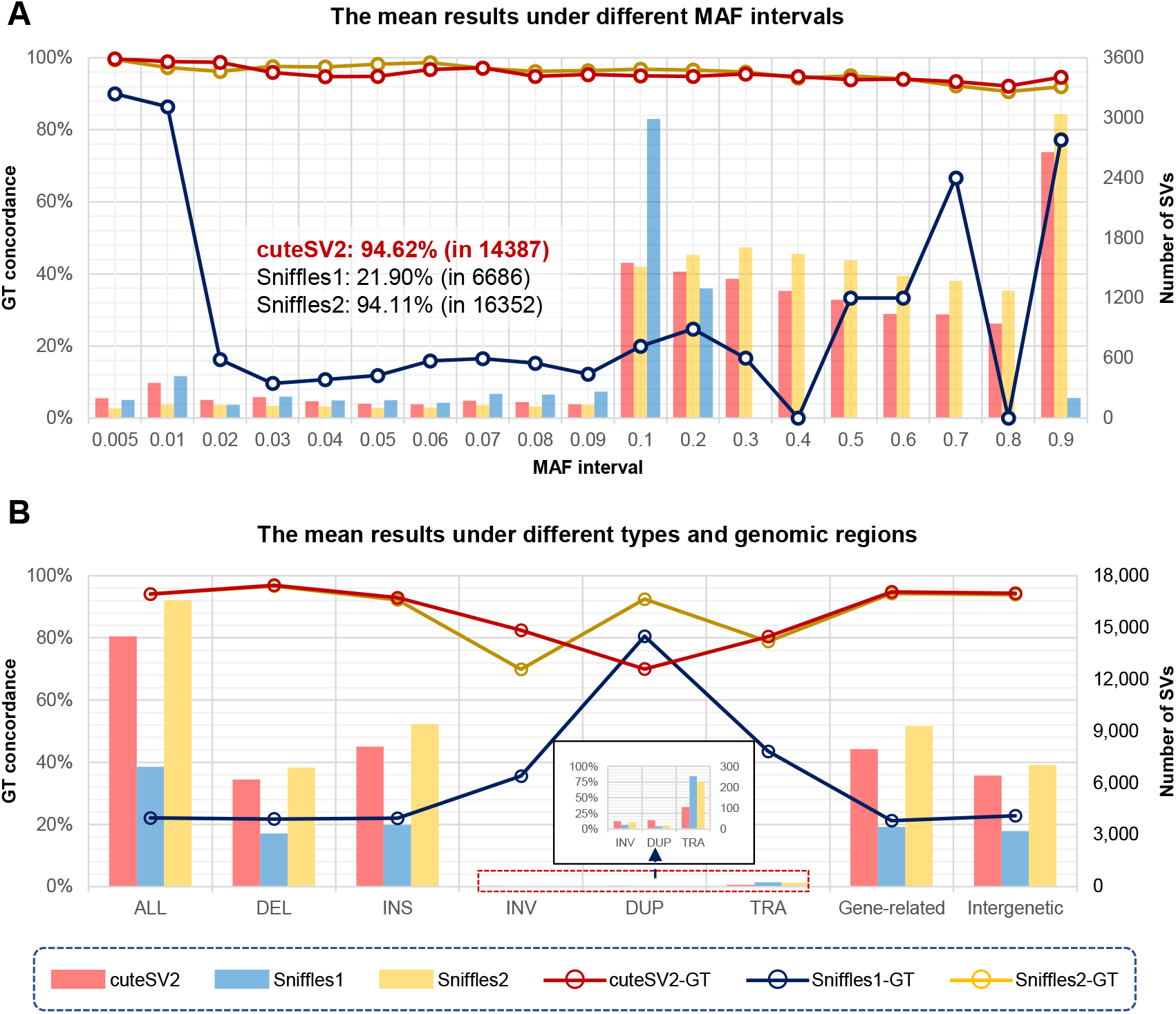
The benchmark results of HiFi validation on the Chinese population group. **A**. The average distribution of genotype concordance rates and SV amounts under different minor allele frequency (MAF) intervals for the three selected individuals. **B**. The mean genotype concordance rates and SV amounts for the three tested samples on different SV types (deletion (DEL), insertion (INS), inversion (INV), duplication (DUP), translocation (TRA)) and different genomic regions (gene-related and intergenetic regions). Each column and line represent the mean number of SVs and mean genotype concordance rate of the three samples, respectively.

Furthermore, we studied the performance of SV genotyping of various SV types. Fig. 6B indicates that for insertions and deletions, the conclusion is similar to that above; that is, the highest genotype concordance was obtained using cuteSV2, followed closely by Sniffles2. The most insertions and deletions (9,412 insertions and 6,890 deletions) were obtained using Sniffles2, fewer insertions (8,105 insertions and 6,202 deletions) were obtained using cuteSV2 and fewer insertions and deletions (3,571 insertions and 3,080 deletions) were obtained using Sniffles1. Approximately 30 inversions for each sample were retained in each method, and the inversion genotype concordance of cuteSV2 and Sniffles2 was apparently higher than that of Sniffles1. In regard to duplications, approximately twice as many duplications were retained using cuteSV2 as the other two methods, and the genotype concordance did not vary much among the different methods (Supplementary Table S15). Hence, cuteSV2 not only outperformed the other two methods with respect to the number of insertions and deletions but also in obtaining longer and more complex SVs.

We also took the various genomic regions where SVs were involved into account. It is obvious that cuteSV2 achieved almost the highest genotype concordance for those SVs located in gene-related regions (Fig. 6B and Supplementary Table S16). This result indicates that cuteSV2 had a highly accurate SV detection ability, especially in high genetic conservation regions, which would benefit studies that rely heavily on functional SVs.

### 4. Evaluation of the computational performance

Finally, we examined the computational performance of these tools on different datasets. From the assessments of elapsed time on the HG002 sample (Fig. 7A and Supplementary Table S17), Sniffles2 had the fastest speed regardless of the dataset (i.e., HiFi: 790 s, CLR: 7,899 s, ONT: 4,587 s), with cuteSV2 (i.e., HiFi: 1,317 s, CLR: 19,028 s, ONT: 12,946 s) following at approximately one time slower, while Sniffles1 (i.e., HiFi: 5,587 s, CLR: 24,525 s, ONT: 16,742 s) and SVJedi (i.e., HiFi: 110,464 s, CLR: 106,683 s, ONT: 122,960 s, all under 16 threads) were much slower. Notably, a quasilinear speedup and a time reduction with increasing threads were observed for cuteSV2 and Sniffles2. Hence, there would be no significant time discrepancy between cuteSV2 and Sniffles2 when applying more CPU threads.

**Fig. 7.**
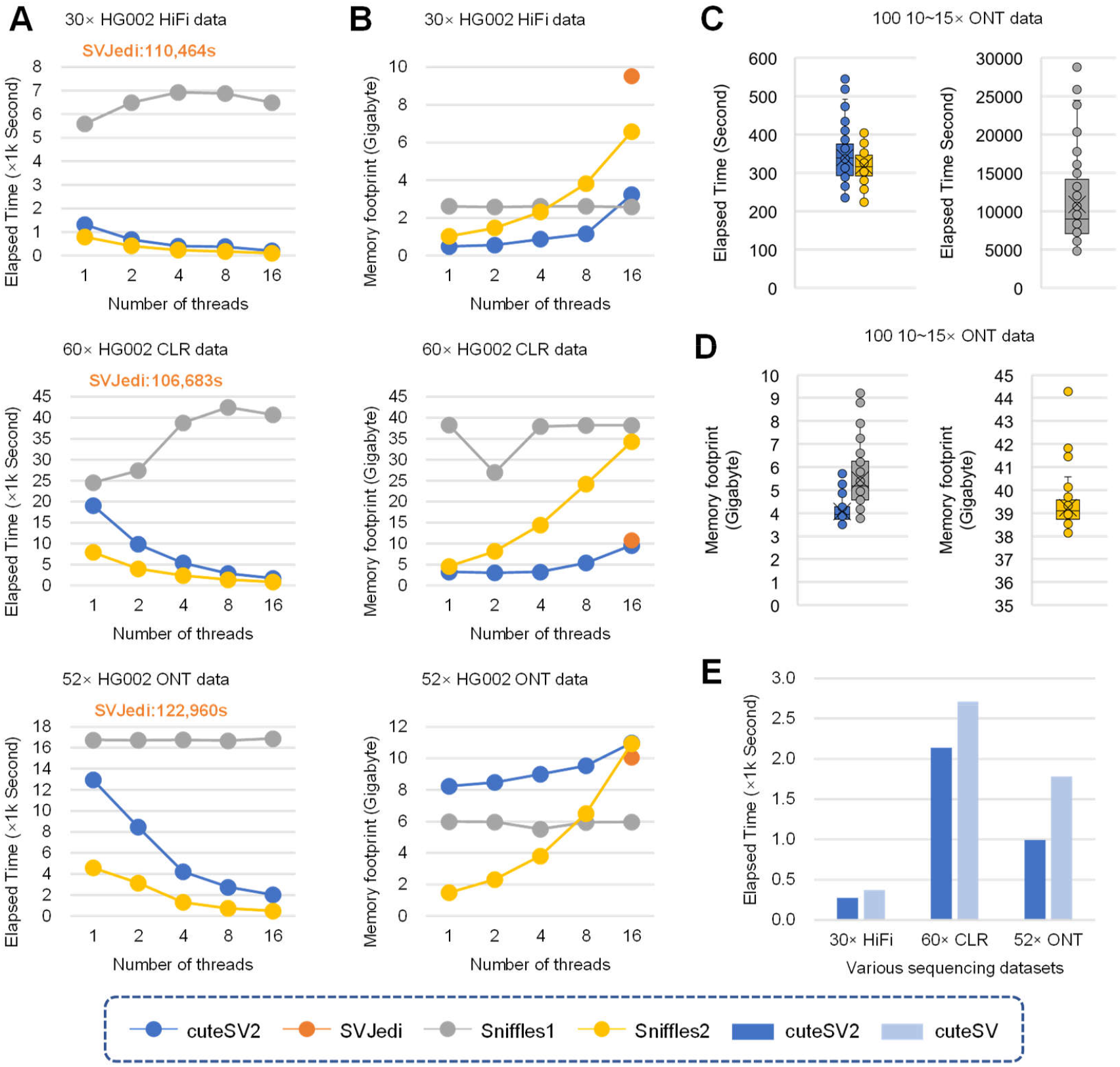
The computational performance for each tool. **A**. The elapsed time on the different HG002 datasets (PacBio HiFi, PacBio CLR, and ONT) with various threads (1, 2, 4, 8, 16). **B**. The memory footprint on the different HG002 datasets (PacBio HiFi, PacBio CLR, and ONT) with various threads (1, 2, 4, 8, 16). **C**. Box plot of the elapsed time of the 100 Chinese individuals. **D**. Box plot of the memory footprint of the 100 Chinese individuals. **E**. The elapsed time of cuteSV2 and origin cuteSV on the different HG002 datasets (PacBio HiFi, PacBio CLR, and ONT) with 16 threads.

In terms of memory footprint, an increasing memory footprint was observed with multiple threads when using cuteSV2 and Sniffles2. This increase was gentler for cuteSV2 than for Sniffles2, whereas a more stable but larger memory footprint was observed using Sniffles1 (Fig. 7B). On the HiFi dataset, only 0.49 GB was utilized under a single thread in cuteSV2. When increasing to 16 threads, smaller memory footprints were observed for cuteSV2 and Sniffles1 (3.32 GB for cuteSV2, 2.58 GB for Sniffles1), whereas a higher memory cost was noted for Sniffles2 and SVJedi (6.57 GB for Sniffles2, 9.51 GB for SVJedi). On the CLR datasets, the fewest memory footprints were expended for cuteSV2 and SVJedi, and much larger footprints were noted for Sniffles1 and Sniffles2. On the ONT datasets, Sniffles1 performed much better than the other three methods, with only a 5 GB memory footprint under various threads. The other three methods cost approximately 10 GB memory footprints for 16 threads.

For the assessments of the data from 100 Chinese individuals (Fig. 7C, 7D and Supplementary Table S18), equivalent and fast speeds were achieved using cuteSV2 and Sniffles2 (i.e., 340 s and 319 s per sample on average, respectively), whereas the average memory footprint for cuteSV2 (4.06 GB per sample) was approximately ten times smaller than that for Sniffles2 (39.23 GB per sample). Sniffles1 required significantly more time (10,884 s) and memory (5.44 GB) on each sample. In conclusion, the least number of computational resources on the regenotyping of a large cohort were expended using cuteSV2.

In addition, in cuteSV2, the SV discovery calling module of cuteSV was upgraded, resulting in the efficient acceleration of the genotyping module in SV discovery. We compared the elapsed time to detect SVs in the HG002 datasets under various sequencing technologies (enabling the genotyping module) (Supplementary Table S19). Fig. 7E demonstrates that there was a large time reduction for the discovery under all sequencing technologies, especially on the ONT datasets, and the time elapsed was reduced from 1780 s to 992 s, which was about twice as fast. On the HiFi and CLR datasets, the time elapsed also dropped more than that of the origin cuteSV (HiFi: from 372 s to 276 s, CLR: from 2,707 s to 2,136 s).

## Discussion

Through regenotyping SVs via long reads, we introduced an accurate and fast regenotyping solution using specifically designed algorithms that effectively promotes studies, such as population genetic measurements, and further analysis. To better apply cuteSV2 in practice, there are still several advantages and disadvantages that need to be explained in detail.

- A two-round purification on SV signatures is implemented in cuteSV2 to select the reads that support the target SV. In the first round, the refinement step of cuteSV is followed to distinguish signatures with various allele lengths, while in the second round, an enhanced purification strategy is performed by comprehensively inspecting the allele similarity. In the example shown in Fig. S1, a target deletion (1,620 bp) on chromosome 13 was correctly obtained by using cuteSV2, and the correct genotype was reported. This is mainly due to the strength of the two-round purification strategy for determining the signatures on complex genomic regions.
- In cuteSV2, a merging strategy is applied for fragile signatures affected by the erroneous read alignment, and agglomerated signatures are generated. This will improve the sensitivity of regenotyping through the maintenance of as many potential signatures as possible. In the example shown in Fig. S2, a target insertion (299 bp) on chromosome 1 was determined to be homozygous only when using cuteSV2 and is concordant with the corresponding ground truth. Given the absence or limitation of the merging strategy, incorrect genotypes were reported using Sniffles1 and Sniffles2.
- The heuristic SV signature extraction module enables cuteSV2 to discover a larger number of signatures and more accurate signatures, especially for longer and more complex SVs such as inversions and duplications, which helps to carry out high-quality genotype assignments. In Fig. S3, there is an inversion of 1,200 bp on chromosome 1, and the real genotype was only successfully reported using cuteSV2. In another example, there is a duplication of 7,759 bp on chromosome 1, as shown in Fig. S4. The genotype is reported as 0/0 in all methods except cuteSV2. This duplication is located on the *HRNR* gene, which is highly associated with gene compression, and the correct genotype assignment is only achieved using cuteSV2, which indicates that cuteSV2 has the potential for functional SV discovery in accurate clinical studies.
- Accurate regenotyping also benefits SV analysis in a large cohort. Fig. S5 shows a 167 bp insertion on chromosome 14 that widely exists in the 100 Chinese individuals. From the figure, the genotypes of the two individuals (Samples #31 and #63) should be “1/1”. The correct genotype is output using cuteSV2, while outputs of “0/1” and “0/1” are obtained using Sniffles1, and outputs of “0/0” and “0/1” are obtained using Sniffles2. Hence, a higher allele frequency of this insertion of 0.995 is reported using cuteSV2, which is beneficial for the successful detection of widely shared common SVs among the population group.
- The fast speed and low memory expense of cuteSV2 enables its wide application in large-cohort SV studies. When applying regenotyping on 100 individuals in 16 usual threads, the fewest computational resources are utilized by cuteSV2. The other methods, such as Sniffles1, has a similar memory footprint but is over 30 times slower, and Sniffles2 requires a similar CPU time on each sample but needs approximately 10 times more computational memory. Therefore, given the restriction of limited computational resources, cuteSV2 can take full advantage of the resources to apply regenotyping and outperform other methods in the force calling of large-scale populations.

Although cuteSV2 has proven its great ability of regenotyping, there are still several limitations both for it and the current regenotyping technologies.

- On the one hand, cuteSV2 can handle most SV types, including insertions, deletions, inversions, duplications, and translocations. However, the regenotyping of copy number variation (CNV) has been difficult to achieve until now. The undiscovered number of copies introduces complexity when genotyping CNVs. To generate accurate genotypes of CNVs in the future, support from more abundant signatures, such as read depths or consensus, is vital.
- On the other hand, in cuteSV2 and other methods, the aim is to complete regenotyping on the diploid genome but not on the polyploid genome, and this is also a bottom-neck in the variant calling field. For polyploids that consist of multiple haplotypes, it is difficult to ensure the exact haplotype where the variation occurs. Hence, new genotyping modules should be designed for polyploids in the future, which might better resolve the haplotype through local haplotype construction on the target region or assign the genotype information through haplotype phasing. We will further focus on these topics and try to solve them in the future.

## Conclusions

In this article, we introduce an accurate and fast regenotyping method, cuteSV2, to assign genotypes for population-based SVs via long-read sequencing. Due to the two algorithms of heuristic signature purification and specifically designed scanning lines, accurate signature marking and effective read distribution statistics, benefitting the reliable likelihood estimation of SV genotypes, are achieved using cuteSV2. The benchmarking results indicate that cuteSV2 retains outstanding regenotyping performance compared with the state-of-the-art methods in various aspects. In particular, it has high accuracy with all sequencing technologies regardless of the sequencing depth. Its application in a large-scale population cohort also proves that the genotypes assigned via cuteSV2 have better distribution concordance to the known genetic regulation and well-studied worldwide cohort. These strengths demonstrate that cuteSV2 has a reliable genotype recognition ability, contributing to high-quality SV detection for large-scale population studies and precise clinical practice. We believe that cuteSV2 will have wide application in cutting-edge genomic research.

## Methods

The force-calling regenotyping method is an expansion module that was integrated into our previous SV detector cuteSV^[21, 22]^. It can be used to assign new genotypes that refer to the given population-scale SVs in parallel for each SV type on each chromosome among every individual. This approach has four major steps, which are described as follows (Fig. 1).

### Extract signatures of various SV types from alignments

We follow the signature extraction module of cuteSV to comprehensively collect various types of SV signatures, including insertions, deletions, inversions, duplications, and translocations.

### Mark candidate signatures via spatial and allele similarity estimation

For each regenotyped SV call in cuteSV2, all candidate signatures extracted above are marked through specifically designed spatial similarity measurements and allele similarity estimation. First, a binary search strategy is implemented on the ordered signature list to find the signature *Sig*_*flag*_ that has the nearest coordinate as the given SV. Then, the spatial similarity of upstream and downstream signatures is measured and collected recursively. For a collected signature *Sig*_*NN*_, its nearest neighborhood signature *Sig*_*NN*_′ will be collected when it satisfies:

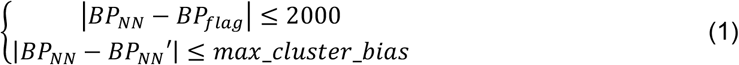

where *BP*_*NN*_, *BP*_*NN*_′ and *BP*_*flag*_ are the breakpoints of *Sig*_*NN*_, *Sig*_*NN*_′ and *Sig*_*flag*_, respectively; *max_cluster_bias* indicates a threshold that is mutative due to various SV types and sequencing platforms (by default, 200, 100, 500, 500 and 50 for DEL, INS, DUP, INV and TRA, respectively). It is worth noting that for translocations, an additional condition that constrains the same transferred chromosome ID of signatures is designed to ensure the spatial similarity of collected signatures.

In the above-collected signatures, only the spatial similarity is considered, while the similarity in alleles is ignored. Therefore, the collected signatures are then purified through allele similarity estimation, and only signatures with high allele similarity are retained as candidate signatures. For insertions and deletions, the fragile signatures from the same read are merged to generate novel potential signatures to restore as many signatures as possible. All these signatures are sorted by their variation length. Then, a quasi-Markov model-based clustering strategy is applied to the refinement step of cuteSV. The *i*th signature is in the same cluster with the *i* − 1th signature if their difference in length is smaller than a dynamic *threshold*_*i*_ defined in (2).

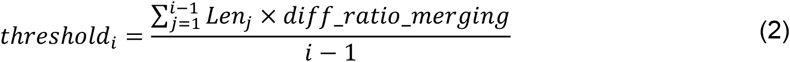

where *diff_rati0_merging* represents a ratio that is mutative due to various SV types and sequencing platforms. After clustering, and the allele similarity between each cluster and the given regenotyped SV is measured as follows:

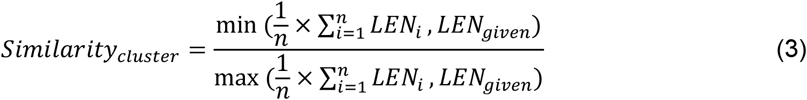

where *LEN*_*given*_ and *LEN*_*i*_ indicate the length of the given regenotyped SV and the *i* th signature in the cluster, respectively. The cluster with the highest allele similarity is selected as the candidate signatures. For inversions, duplications, and translocations, the collected signatures are used directly.

### Mark overlapped reads for read distributions

The distribution of reads around each regenotyped SV breakpoint is computed using an overlapping scanning line. Specifically, for each SV, all alignment reads that overlap the SV on the chromosome are recorded.

All the regenotyped SVs and alignment reads are collected as a 2-tuple (*s*_*SV*_, *e*_*SV*_), (*s*_*read*_, *e*_*read*_), where *s*_*read*_ and *s*_*SV*_ represent the start coordinate of read/SV, *e*_*read*_ and *e*_*SV*_ represent the end coordinate of read/SV, respectively. The tuples are grouped by chromosome, and in each group, all the breakpoints in the tuples are sorted in order according to their coordinates. Then, a line scanning the ordered breakpoints from head to tail is designed. While moving the scanning line, a read set *R* is used to record the reads that stride over the line and *R* + {*read*|*s*_*read*_ < *bp*_*line*_ < *e*_*read*_}, where *bp*_*line*_ represents the breakpoint that the scanning line reaches. In detail, the read is added into the read set when the scanning line reaches *s*_*read*_ and removed from the read set when the line reaches *e*_*read*_. When the line reaches *s*_*SV*_ or *e*_*SV*_, reads in read set *R* are recorded as the reads covering the SV breakpoint. The intersection of reads covering both start and end breakpoints of an SV is marked as the reads that overlap the SV.

### Assign genotypes based on likelihood estimation

The reads of candidate signatures are subtracted from the overlapped reads to retain the read set that matches the reference. Using this read set according to the reference and the candidate signatures according to the allele, the genotyping module from cuteSV is utilized to assign genotypes through the biallelic assumption on the likelihood of various zygosities.

### Process the benchmarking datasets

For the Ashkenazim Trio, we obtained the PacBio HiFi sequencing alignment data of HG002, HG003, and HG004 from GiaB and applied PBSV (v 2.4.0) to detect the individual SVs. The cohort-level structural variations were generated by merging the three individuals using SURVIVOR^[23]^ (v 1.0.7). Then, we obtained CLR and ONT sequencing alignment data of HG002 from GiaB and regenotyped the cohort-level structural variations using four methods (cuteSV2 (v 2.0.2), Sniffles1 (v 1.0.12), Sniffles2 (v 2.0.2), SVJedi (v 1.1.6)).

For the 100 samples from the Chinese population group, we first implemented ONT sequencing. Four different callers (cuteSV (v 1.0.13), SVIM^[24]^ (v 2.0.0), Sniffles1, and Nanovar^[25]^ (v1.3.9)) were applied for individual SV calling, and SURVIVOR was applied for sample-level merging through various callers. After obtaining 100 sample VCFs (each one was merged from four callers), they were integrated using SURVIVOR a second time to generate cohort-level population VCFs. With the merged cohort-level SVs, we implemented regenotyping methods on 100 samples and applied SURVIVOR a third time to obtain the revised cohort-level SV callsets. Then, we implemented PacBio HiFi sequencing on three of these samples (i.e., Samples #1, #2, and #3) and applied PBSV to these three HiFi samples to achieve reliable sample SVs.

The detailed commands can be found in the Supplementary Note.

### Evaluate the regenotyping performance

The regenotyping results were benchmarked using Truvari^[26]^ (v 3.2.0); more details of various aspects are provided below. The precision, recall, and F1 score are defined as follows:

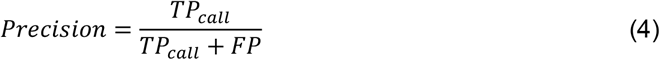

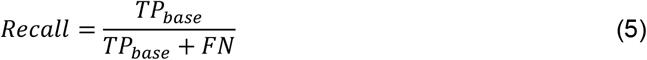

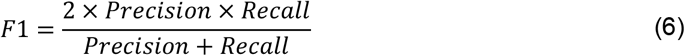

where *TP* _*call*_ and *FP* represent the number of SVs that are concordant and inconcordant with the ground truth, respectively, and *TP*_*base*_ and *FN* represent the number of SVs in the ground truth that are discovered and missing, respectively. Genotype concordance is defined as

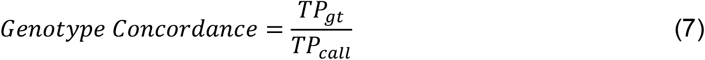

where *TP*_*gt*_ represents the number of SVs with concordant genotypes to the ground truth. Furthermore, the tests of HWE and ExcHet were achieved using the bcftools (v 1.9)^[20]^ fill-tags module.

The benchmarks were implemented using a server with 2 Intel(R) Xeon(R) Gold 6240 CPUs @ 2.60 GHz (32 cores in total), 15.8 terabytes of RAM and 6221 terabytes hard disk space (no SSD was used), running on the CentOS Linux release 7.5.1804 operating system. The elapsed time and memory footprint were assessed by using the “seff” command of the Slurm Workload Manager.

## Declarations

### Ethics approval and consent to participate

Not applicable

### Consent for publication

Not applicable

### Code availability

The force calling module in cuteSV2 was implemented in Python and can be easily installed via bioconda and PyPI. Its source code is available at https://github.com/tjiangHIT/cuteSV^[27]^. The cuteSV2 release used in this article is deposited on Zenodo with doi: https://doi.org/10.5281/zenodo.7304294^[28]^.

### Data availability

The alignment data in the Ashkenazim Trio experiment are available at https://ftp-trace.ncbi.nlm.nih.gov/ReferenceSamples/giab/data/AshkenazimTrio/. The original data of 100 Chinese individuals are restricted; please contact for permission to acquire these data.

## Author contributions

TJ, SC, YL and BL designed the method. SC and TJ implemented the method. SC and SL performed the experiments and data analysis. SC and TJ wrote the manuscript. The authors read and approved the final manuscript.

