## Supplementary Figures and Notes for "Regenotyping structural variants through an accurate force-calling method"

**Supplementary Material**

Tao Jiang^1,†,*^, Shuqi Cao^1,†^, Yadong Liu^1^, Shiqi Liu^1^, Bo Liu^1^, GuoHua Wang^1,*^ and Yadong Wang^1,2,*^

^1^Faculty of Computing, Harbin Institute of Technology, Harbin 150001, China.

^2^School of Medicine and Health, Harbin Institute of Technology, Harbin 150001, China.

^†^Joint first authors: Tao Jiang and Shuqi Cao.

**Content**

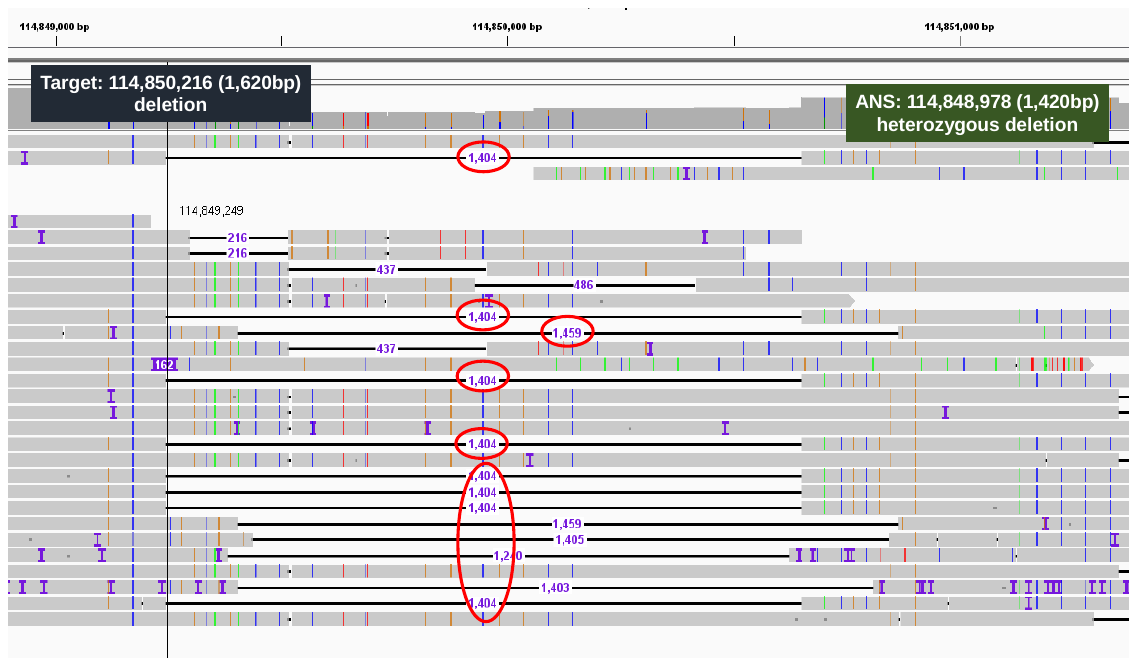

**Fig. S1. An example of a deletion only being detected by cuteSV2 on the 30× HG002 via PacBio HiFi data**

According to the ground truth, there is a 1,404 bp heterozygous deletion at chr13: 114,848,978, and the target SV is 1,602 bp at chr13: 114,850,216. From the IGV snapshot, there are many insertion signatures at a similar location around this event and they have various lengths (from 400 bp to 1,400 bp). Sniffles1 reported a “1/1”, and Sniffles2 and SVJedi gave a “0/0” record. By contrast, cuteSV2 purified these signatures and only remained 13 signatures supporting the deletion which are marked by the red circle in the figure. With these correct supporting reads, only cuteSV2 outputted the genotype “0/1” for this deletion correctly.

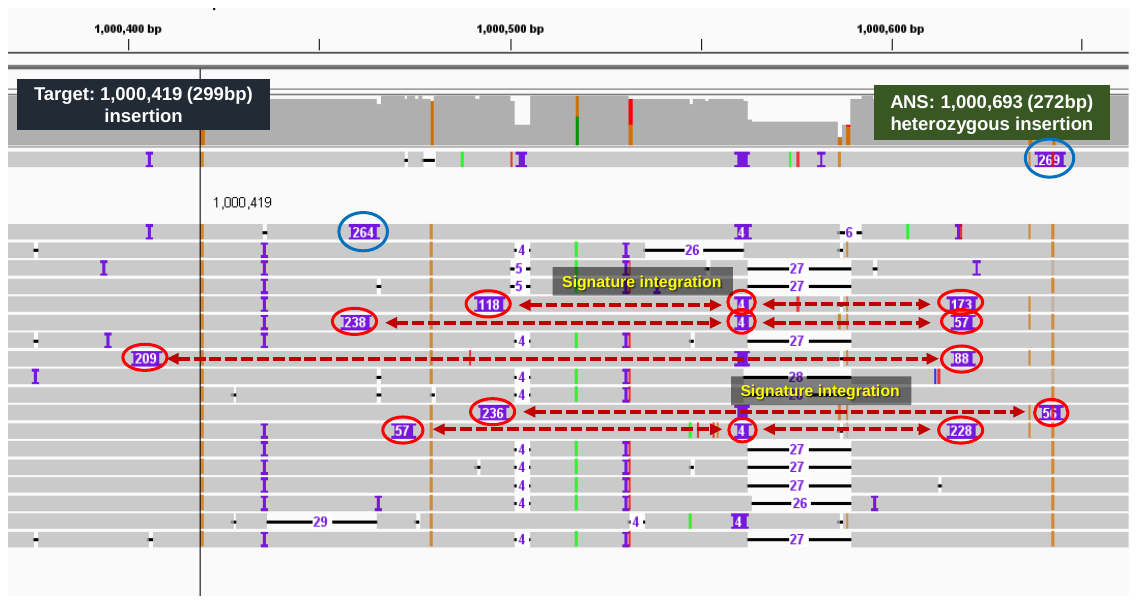

**Fig. S2. An example of an insertion only being detected by cuteSV2 on the 30× HG002 via PacBio HiFi data**

According to the ground truth, there is a 272 bp heterozygous insertion at chr1: 1,000,693, and the target SV is 299 bp at chr1: 1,000,419. The IGV snapshot shown above indicates that there are many fragile insertion signatures with various lengths (from 50 bp to 300 bp) around this event. A few fragments have a similar length to the target SV (marked in blue circles), while the other fragments appear in multiple places of the same read and the addition of their length corresponds to the target SV (marked in the red circles). In this case, Sniffles1 and SVJedi reported it as “1/1”, and Sniffles2 gave a “0/0” record. However, a signature integration on these fragile signatures was implemented by cuteSV2 to generate potential signatures and recovered most insertion signatures. In this way, cuteSV2 finally outputted the genotype “0/1” for this insertion correctly.

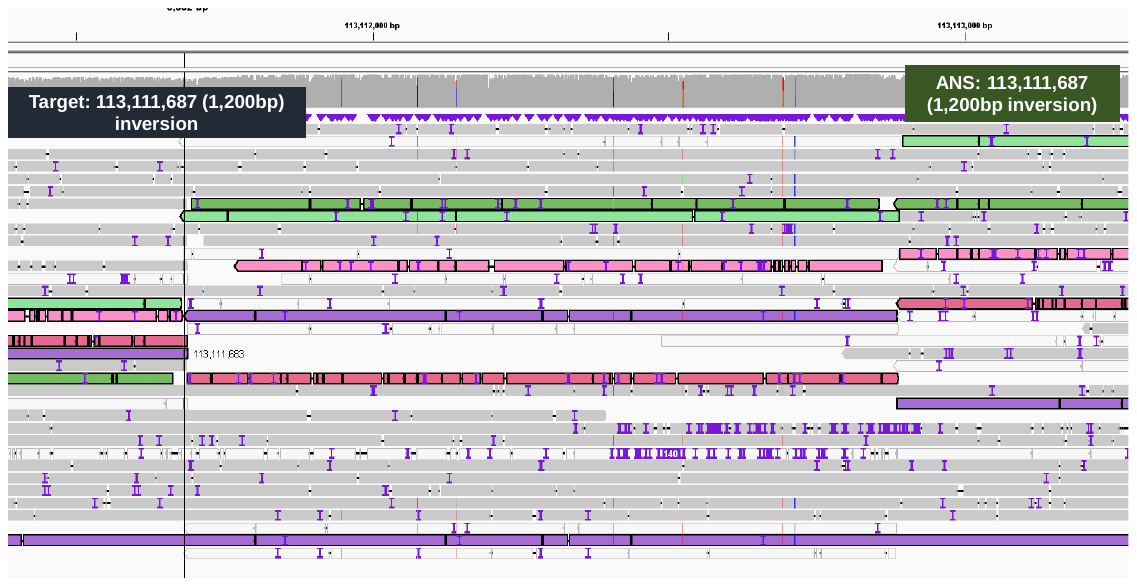

**Fig. S3. An example of an inversion only being detected by cuteSV2 on the 15× Sample #2 via ONT PromethION data**

According to the ground truth, there is a 1,200 bp heterozygous inversion at chr9: 113,111,687, and the target SV is 1,200 bp at chr9: 113,111,687. The IGV snapshot of the shown figure also proves the occurrence of the inversion due to the inverse supplementary alignments which are marked by different colors. However, Sniffles1 and Sniffles2 both gave “0/0” records for this inversion. Based on the various signature extraction from cuteSV2, it first fully discovered the inverse supplementary alignment segments, and then analyzed the distribution of the reads around them and reported this heterozygous inversion successfully.

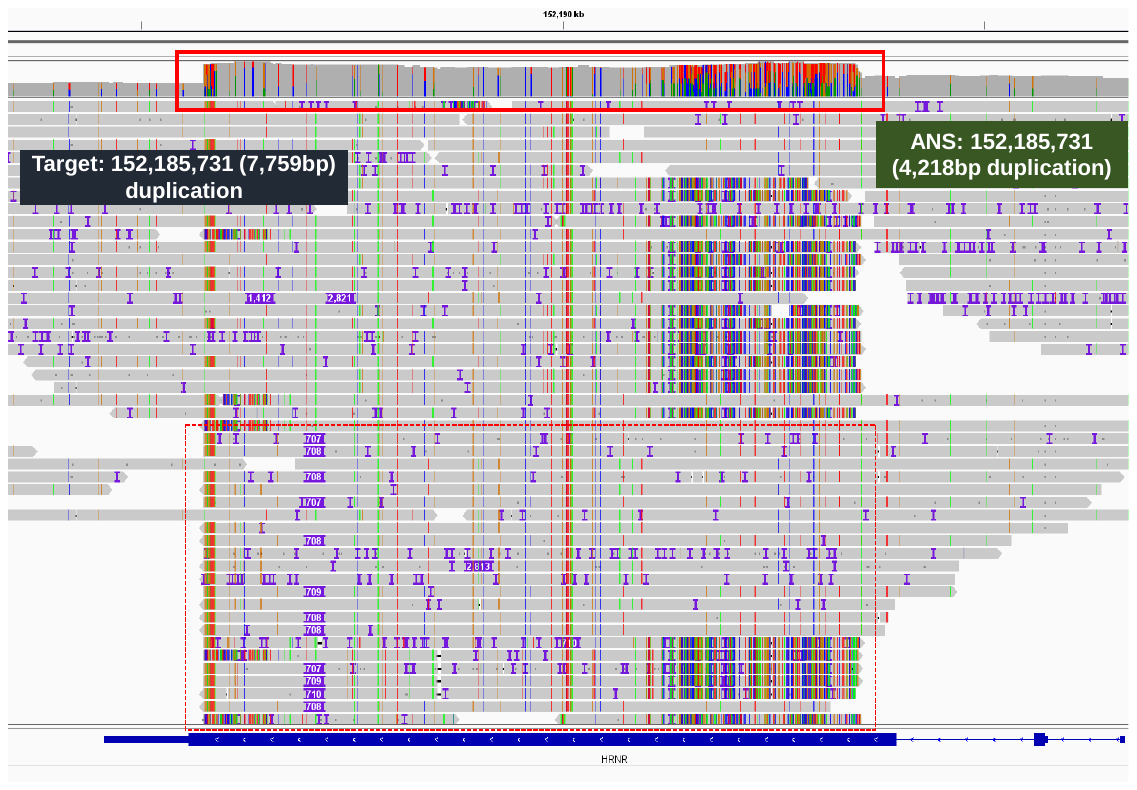

**Fig. S4. An example of a duplication only being detected by cuteSV2 on the 30× HG002 via PacBio HiFi data**

According to the ground truth, there is a 4,218 bp heterozygous duplication at chr1: 152,185,729, and the target duplication of 7,759 bp at chr1: 152,185,731. From the IGV snapshot, it is obvious that there exists a duplication due to the local increasing coverage compared to the nearby region. Sniffles1 and Sniffles2 regenotyped this duplication as “0/0”. By contrast, based on the various signature extraction from cuteSV2, cuteSV2 collects the complex corresponding reads and extracts the duplication signatures, which helps to detect the duplication successfully. Furthermore, we find the duplication is located in the *HRNR* Gene, which has been proven associated with many diseases including Spastic Paraplegia, Autosomal Recessive, and Ichthyosis Vulgaris. Therefore, this duplication related to the gene region would probably influence the gene expression. The successful detection and regenotyping of it for cuteSV2 would bring potential in clinical diagnosis and some studies related to gene analysis.

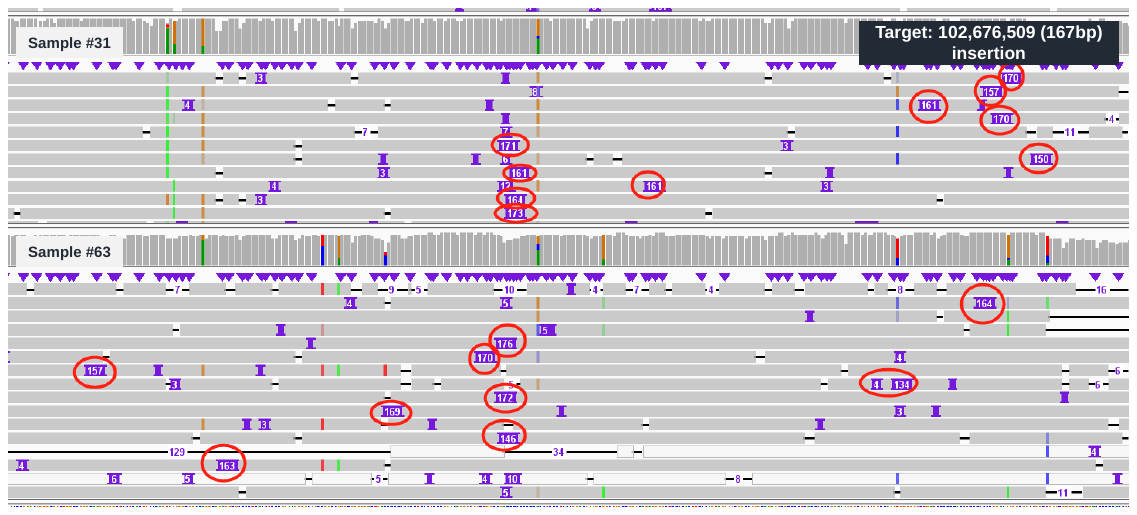

**Fig. S5. An example of an insertion only being reported with correct allele frequency by cuteSV2 on the population cohort via ONT PromethION data**

According to the IGV snapshot, there is a homozygous 167bp insertion around chr14:102,676,509. Also, this insertion widely exists among all the 100 Chinese individuals as a variation of high allele frequency. However, Sniffles1 and Sniffles2 gave “0/1”, “0/1” and “0/0”, “0/1” records for this insertion, respectively. cuteSV2 collected the abundant SV signatures and genotyped both samples as “1/1” records. Hence, only cuteSV2 reported a higher allele frequency insertion here (AF: 0.995 for cuteSV2, 0.965 for Sniffles2, and 0.5 for Sniffles1).

**Supplementary Notes**

The complete commands of data generation, variant calling and benchmarking implementation in this study are as below:

**1. Implementation of Ashkenazim Trio**

> pbsv discover {long_reads.bam} {long_reads.svsig.gz} --tandem-repeats human_hs37d5.trf.bed

> pbsv call {reference.fa} {long_reads.svsig.gz} {sv_calling.vcf} --ccs -t INS,DEL

> cat {*.sv_calling.vcf} > {merge_files.txt}

> SURVIVOR merge {merge_files.txt} 1000 1 1 0 1 30 {population.vcf}

> samtools view -h -s {ratio} {long_reads.bam} | samtools view -bS > {down_sample.bam}

> samtools index {down_sample.bam}

**2. Implementation of Chinese Population Group**

> bcftools +fill-tags -o {force_calling.filltags.vcf} {force_calling.vcf}

> truvari anno bpovl -i {force_calling.vcf} -a Homo_sapiens.GRCh38.106.gtf.gz -o {force_calling.anno.jl} -p gff

> python gene_anno.py {force_calling.anno.jl} {force_calling.vcf} {gene.vcf} {inter_gene.vcf}

**3. Implementation of force calling**

**cuteSV2 calling**

- **For PacBio CLR data**

> cuteSV {long_reads.bam} {reference.fa} {force_calling.vcf} {workdir} --max_cluster_bias_INS 100 --diff_ratio_merging_INS 0.3 --max_cluster_bias_DEL 200 --diff_ratio_filtering_DEL 0.5 -sl 30 -Ivcf {population.vcf} -q 10

- **For PacBio HiFi data**

> cuteSV {long_reads.bam} {reference.fa} {force_calling.vcf} {workdir} --max_cluster_bias_INS 1000 --diff_ratio_merging_INS 0.9 --max_cluster_bias_DEL 1000 --diff_ratio_filtering_DEL 0.5 -Ivcf {population.vcf} -q 10

- **For ONT data**

> cuteSV {long_reads.bam} {reference.fa} {force_calling.vcf} {workdir} --max_cluster_bias_INS 1000 --diff_ratio_merging_INS 0.3 --max_cluster_bias_DEL 1000 --diff_ratio_filtering_DEL 0.3 -Ivcf {population.vcf} -q 10

**Sniffles1 calling**

> sniffles -m {long_reads.bam} -v {force_calling.vcf} --Ivcf {population.vcf}

**Sniffles2 calling**

> sniffles --input {long_reads.bam} --vcf {force_calling.vcf} --genotype-vcf {population.vcf}

**SVJedi calling**

> samtools fasta {long_reads.bam} > {long_reads.fasta}

> python3 svjedi.py -v {population.vcf} -r {reference.fa} -i {long_reads.fasta} -o {force_calling.vcf}

**4. Assessment on the sensitivity, accuracy, runtimes and memory footprints**

> bgzip -c {force_calling.vcf} > {force_calling.vcf.gz}

> tabix {force_calling.vcf.gz}

> truvari bench -b HG002_SVs_Tier1_v0.6.vcf.gz -c {force_calling.vcf.gz} --includebed HG002_SVs_Tier1_v0.6.bed -o cmp -p 0 -r 1000 --multimatch –passonly

> truvari bench -b HG002_GRCh37_CMRG_SV_v1.00.vcf.gz -c {force_calling.vcf.gz} --includebed HG002_GRCh37_CMRG_SV_v1.00.bed -o cmp -p 0 -r 1000 --multimatch –passonly

> seff {job_id}
